# Constructing microbiome co-occurrence networks with confidence: A conditional, nonparametric, inference-based approach

**DOI:** 10.64898/2026.08.27.747483

**Authors:** Hoseung Song, Yunhua Xiang, Hongjiao Liu, Wodan Ling, Anna M. Plantinga, Sujatha Srinivasan, Yuzheng Dun, Ni Zhao, Shan Sun, Stephanie M. Engel, Noah Simon, Michael C. Wu

## Abstract

Constructing microbial association networks is a common strategy for exploring relationships among taxa in microbiome studies. Although marginal correlation methods are easy to implement and allow formal inference, they can produce spurious edges driven by indirect associations through other taxa. Conditional graphical-modeling methods aim to recover direct associations, but many rely on Gaussian or linear assumptions and often provide limited uncertainty quantification. We propose a conditional, nonparametric approach based on the scaled expected conditional covariance (SEcov). SEcov measures population-level conditional association by residualizing each taxon with respect to the remaining taxa and scaling the resulting expected conditional covariance. The resulting estimator can incorporate flexible machine-learning methods for conditional-mean estimation and admits asymptotic normal inference, enabling p-values and confidence intervals for taxon-pair associations. We demonstrate through simulation studies that our proposed approach improves network recovery relative to other methods, and we illustrate the new method via construction of a co-occurrence network for the vaginal microbiome during pregnancy.

**IMPORTANCE:** High-throughput sequencing has made it possible to characterize microbial communities at large scale, and network analysis is widely used to summarize relationships among taxa. However, networks based on marginal correlations may include indirect associations, whereas many conditional graphical models rely on assumptions that may be difficult to justify for sparse, zero-inflated, compositional microbiome data. SEcov offers a practical alternative by estimating conditional associations nonparametrically and attaching inferential uncertainty to individual edges. This allows investigators to construct microbiome networks using statistically interpretable evidence for taxon-pair associations, rather than relying solely on arbitrary correlation cutoffs or regularization tuning parameters.

## INTRODUCTION

Building co-occurrence networks is a standard analytic procedure in microbiome profiling studies that sheds important light on relationships among bacterial taxa, providing clues as to the mechanisms underlying the impact of microbial communities on host conditions. Construction of networks generally involves creating an undirected network in which nodes represent taxa and edges connecting two adjacent nodes represent the association and dependency relationship between them. Intuitively, a greater magnitude of the association measure implies a higher chance of co-occurrence, suggesting greater interaction between the two taxa (nodes) within the community. Consideration of all pairs of taxa allows full characterization of the interactions among all involved taxa. Such networks have been utilized to identify key factors sharing a common role in an ecosystem, such as soil microbial communities [3], carbon flux in the oceans [14], and nitrogen-transforming networks [20].

Given interest in construction of co-occurrence networks, a range of computational and statistical approaches have been developed. Generally, these approaches fall into one of two classes, unconditional correlation analysis and conditional correlation analysis. These two strategies have distinct conceptual and operational characteristics.

The idea underlying unconditional correlation analysis is to assess the relationship between each pair of taxa by assessing their correlation, one pair at a time, without consideration for the other taxa in the community. Operationally, this can be done using Pearson’s correlation or using more sophisticated modeling approaches. The magnitude of the correlation then represents the edge-weight and the strength of the relationship between the pair of taxa. A major strength of this strategy is ease of implementation and formal inference, i.e., computation of confidence intervals and *p*-values for assessing statistical significance of edges. However, the detected associations for unconditional correlation analysis may be problematic due to the *indirect effects* [4]. For example, two taxa called “interacting” may actually be mechanistically associated with the third one but are otherwise unrelated. The correlation between the two taxa is merely a result of failure to condition on their mutual relationships with the third taxon. Consequently, such co-occurrence networks are generally over-complicated with too many spurious connections leading to difficulties in interpretation. Specific unconditional correlation analysis methods include SparCC [12] among others. In contrast to unconditional correlation methods, conditional correlation methods focus on assessing the relationship between pairs of taxa conditioning on all other taxa in the community, primarily through Gaussian graphical models. By conditioning and fixing the abundance of all other taxa, conditional approaches preclude the existence of spurious correlations that result from indirect connections through the other community members. Thus, all correlations are, in principle, direct relationships between taxa. Most strategies for conditional analysis utilize partial correlation [2, 6] in Gaussian graphical models, adjusting for the linear effects of all the other taxa to assess of the linear association between the pair of interest. When data is drawn from a Gaussian distribution, this partial correlation can work well since the underlying dependency structure is completely characterized by linear conditional dependence [11, 37]. Due to this fact, the network can be fully encoded by the inverse covariance matrix, also called the precision matrix: two variables with non-zero entries in the precision matrix are conditionally dependent and are connected by an edge. This corresponds to assessing the conditional dependency in Gaussian graphical models [37]. Many recently developed methods for inferring microbial networks apply this idea, such SpiecEasi [19] among many others [9, 16, 36].

However, despite the interpretability and calculation of a more meaningful quantity, conditional network construction methods that use Gaussian graphical models suffer from other challenges, including very strong parametric (normality and linearity) assumptions and difficulties in formal statistical inference. For example, the conditional dependency and interpretation of models from SpiecEasi and related approaches rely entirely on the Gaussian assumption. Failure to satisfy normality negates the interpretation as the precision matrix or the partial correlation may not fully capture the conditional dependency and thus edges in the detected measures can be spurious. Approaches that try to relax the normality assumption still make strong parametric assumptions. Similarly, most approaches assume linear relationships between pairs of taxa. However, such assumptions are often violated in practice, as data transformations required for compositional microbiome data, such as log-ratio transformations and pseudo-count adjustments for zero-inflation—frequently introduce severe non-linear artifacts. Finally, even when the normality and linearity assumptions are satisfied, it is challenging to obtain *p*-values and confidence intervals to evaluate whether edges and associations between taxa are truly present or just due to random chance.

In this paper, we propose an alternative approach that shares the strengths of the conditional and unconditional approaches, while not inheriting their weaknesses. Specifically, we consider using a more general form of conditional association measure, the Scaled Expected Conditional Covariance (SEcov), for recovering microbial co-occurrence networks. This measure addresses several important issues in the existing works of network analysis. First of all, SEcov is defined without specifying a parametric joint distribution for the data or a particular linear graphical structure among OTUs. Through flexible conditional-mean estimation, it can adjust for nonlinear effects of the conditioning nodes and reduces to partial correlation under the Gaussian assumption. Thus, SEcov targets a broader notion of conditional association than one encoded solely by the precision matrix. Second, unlike partial correlation or any other local quantities like conditional correlation, SEcov provides a global measure of association, which estimate the conditional association at the population level and will not vary with the abundance of OTUs. As a result, one would derive a much more robust microbial network. In addition, it does not require recovery of the precision matrix and can easily accommodate for high-dimensional settings. Later, we will see that there exists a natural and theoretically optimal estimator, which enables the leverage of flexible machine learning methods. More importantly, it enables the construction of confidence intervals, which however can be extremely difficult for the existing graphical modeling approaches. This theoretical property greatly enhances the interpretability of the results: the estimated network is no longer determined by selecting any ad hoc cutoffs of Pearson’s correlation or by tuning regularization parameters as in graphical lasso. With SEcov, one can describe the level of statistical confidence in the existence of OTU-OTU associations.

## MATERIALS AND METHODS

Unlike many graphical modeling approaches demanding the normality, SEcov is a nonparametric graphical modeling technique. In this section, we shall first introduce the form of the population parameter for assessing the conditional dependency in general, as well as a simple and natural estimation procedure that leverages the cutting-edge machine learning techniques. Then, we demonstrate how to apply SEcov for microbiome data.

### Estimation and inference of SEcov by flexible machine learning techniques

In a general multivariate problem, we evaluate and estimate the interplay between *d* variables by an undirected graph *G* = (*V, E, P*). *V* represents a set of nodes *{v*_*j*_*}*_1≤*j*≤*d*_, (i.e., *d* variables). *E* represents a set of edges *{e*_*jk*_*}*_1≤*j≠k*≤*d*_: if node *v*_*i*_ and *v*_*j*_ are conditionally dependent, then *e*_*jk*_ = 1; otherwise *e*_*jk*_ = 0. *P* is a collection of weights *{ρ*_*jk*_*}*_1≤*j*≠*k*≤*d*_ expressing the strength and sign of each edge: (i) *ρ*_*jk*_ *>* 0 (*ρ*_*jk*_ *<* 0) represents that *v*_*i*_ and *v*_*j*_ are positively (negatively) associated, *ρ*_*jk*_ = 0 corresponds to *e*_*jk*_ = 0; (ii) a large |*ρ*_*jk*_| implies a strong correlation. Hence, the goal in the context of graphical modeling is to estimate *P* which determines the structure of the network to be recovered. In the existing literature, *ρ*_*jk*_is commonly estimated by Pearson’s correlation coefficient which induces direct association among variables, or by inverse covariance which accounts for indirect association but requires Gaussian assumption. Here, we introduce the Scaled Expected Conditional Covariance (SEcov) [35] for *ρ*_*jk*_, which characterizes the conditional dependency without any model assumptions. The form of the correlation parameter is:

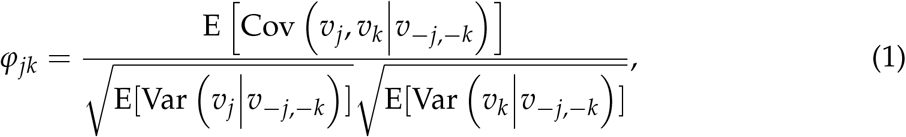

where *v*_−*j*,−*k*_ := *V*/*{v*_*j*_, *v*_*k*_*}* represents the set of all the conditioning variables other than *v*_*j*_ and *v*_*k*_. The construction of *φ*_*jk*_ in (1) is analogous to how Pearson’s correlation is formed from covariance: the numerator includes Cov(*v*_*j*_, *v*_*k*_|*v*_−*j*,−*k*_ ), which is the conditional covariance of *v*_*j*_ and *v*_*k*_ given *v*_−*j*,−*k*_, while the denominator contains two conditional variances, Var(*v*_*j*_|*v*_−*j*,−*k*_ ) and Var(*v*_*k*_|*v*_−*j*,−*k*_ ), serving as the scaling factor. So, *φ*_*jk*_ is scale-invariant and takes on values in [-1, 1] just like Pearson’s correlation does. The difference is in that *φ*_*jk*_ is comprised of local quantities, i.e., the conditional covariance and conditional variance. Because it is adjusted for the effects of the conditioning variables, *ψ*_*j*_*k* is a form of conditional association. In fact, when data is jointly Gaussian, i.e., *V ∼ N*_*d*_(*µ*, Σ), *φ*_*jk*_ reduces to the partial correlation and would be equivalent to methods based on the estimation of precision matrix. However, *φ*_*jk*_ is more general in the sense that its definition does not require joint normality and its estimation can incorporate flexible models for the conditional means, allowing nonlinear effects of the conditioning set to be accommodated. All these facts argue that *φ*_*jk*_ is a useful quantity for recovering the latent conditional dependence structure among all involved variables.

There exists a simple and natural estimator of *φ*_*jk*_ [35]. Suppose that we have *n* observations, 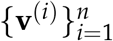 where 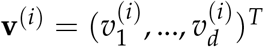 . Then, the conditional association *φ*_*jk*_ between *v*_*j*_ and *v*_*k*_ can be estimated by

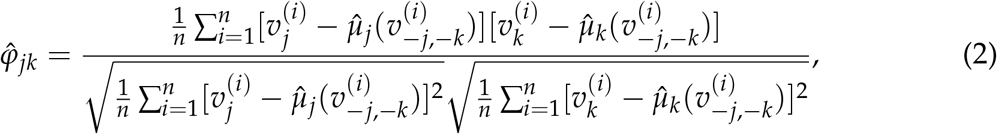

where 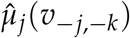 is the estimated conditional mean for the truth *µ*_*j*_(*v*_−*j*,−*k*_ ) := E(*v*_*j*_|*v*_−*j*,−*k*_ ). By (2), we see that the problem of estimating *φ*_*jk*_ is essentially equivalent with estimating two conditional means, *µ*_*j*_(*v*_−*j*,−*k*_ ) and *µ*_*k*_ (*v*_−*j*,−*k*_ ). This is what supervised learning typically does: it builds a model to predict a continuous outcome given a set of features. Hence, the problem of evaluating the conditional dependence is reduced to a *canonical prediction problem*, which allows leverage of many machine learning techniques, such as random forest [21], generalized additive models [15] and so on. These techniques are flexible and are able to capture both linear and non-linear effects of *v*_−*j*,−*k*_, even when the true relationship between *v*_*j*_ (*v*_*k*_) and *v*_−*j*,−*k*_ is unknown and data is high-dimensional. If those conditional means are well estimated, then [35] have shown that 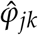 defined in (2) converges to a normal distribution:

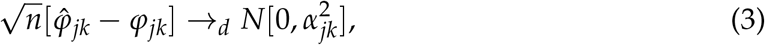

where 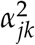 is the asymptotic variance of 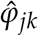 and can be consistently estimated by 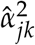 (see Supplemental Material for the form of 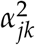 and 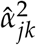 ). The normality shown in (3) implies that 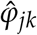 is a centered estimator which converges to a mean-zero normal variable with variance 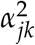 . Due to this nice property, we are then able to address the following hypothesis testing problem:

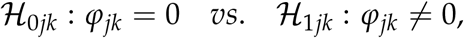

by constructing a test statistic 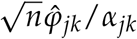 and calculating the *p*-value,

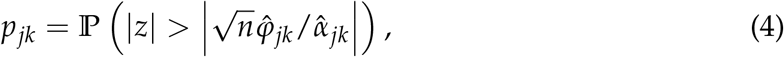

where *z* denotes a standard normal variable. Therefore, under significance level *t*, we only select those edges with *p*_*jk*_ ≤ *t*. This greatly improves the interpretability of the result compared to the existing work. For the Gaussian graphical model or its extensions, one needs to tune a regularization parameter to generate a sparse graph. Methods based on pair-wise correlation also need a pre-specified cutoff and only those edges having correlations greater than the cutoff can be selected. However, neither the regularization parameter nor the cutoff is statistically meaningful. SEcov addresses this issue: choosing a smaller significance level *t* provides stronger statistical evidence that the connected pair of nodes is conditionally dependent..

Next, we shall use SEcov estimator 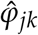 as the basis to assess the OTU-OTU association and construct the undirected network for microbial interactions.

### Preprocessing the compositional data

To apply SEcov to microbiome count data, we first preprocess the data to address two features of amplicon-based sequencing data: unequal library sizes with many zero counts, and the compositional constraint induced by relative-abundance normalization.

Throughout our analysis, the *n* biological samples are treated as independent observations. The compositional issue is different: after counts are converted to relative abundances, the *d* taxon-specific components within each sample are constrained to sum to one. Thus, the taxon proportions cannot vary independently within a sample, and naive covariance, correlation, or regression analyses of relative abundances can produce spurious taxon-taxon associations.

Suppose we have OTU count data for *d* species, collected from *n* individual samples. We denote the count table by a matrix 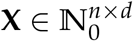 where N_0_ represents the set of non-negative integers. **X** contains *n* row vectors, i.e., 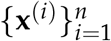 with 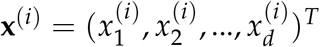, representing the abundance of *d* species in the *i*th sample. Although count-based models can be used to analyze OTU count data directly, raw sequencing counts are affected by sample-specific library sizes and other technical or sampling effects. Therefore, for correlation- or conditional-association-based network estimation, they are not directly comparable across samples without preprocessing. Instead, relative abundance data are often used for analysis and stored in a matrix **Y** with row vectors 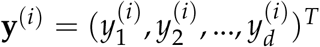, where 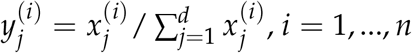 and *j* = 1, …, *d*. This relative abundance normalizes each raw count of a species with respect to the total counts 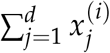 in a sample. This step, however, makes the resulting data compositional: within each sample, the taxon-specific relative abundances are constrained to sum to one. This unit-sum constraint dissuades us from applying those standard statistical tools such as linear regression [28] for low-dimensional data or Lasso [33] for high-dimensional data. Even the analysis of covariance matrix for this compositional data can be biased: a negative correlation trend always exists in **Y** even there is actually no correlation among **X**.

Several methods have been developed to accommodate compositionality. This includes log ratio transformation [1, 17], which exploits the fact 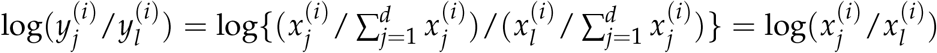, such that any properties possessed by the original abundance ratio also apply to the relative abundance ratio. Often, the adjustment for compositional data is prior to any estimation and inference of the conditional dependence. For example, instead of using the ratio of the counts, [19] uses a centered log-ratio (clr) transformation such that the transformed counts in a sample sum to zero. The clr-transformed count table *V* has the following row vectors:

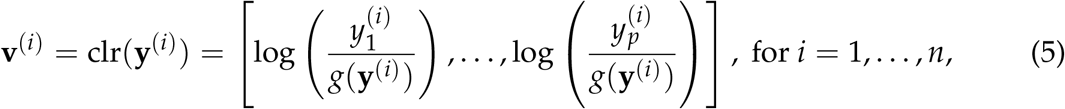

where 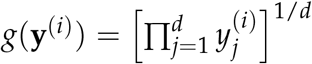 is the geometric mean of each compositional row vector **y**^*(i)*^. This clr transformation also remains symmetric and isometric, i.e., clr(**Y**) = clr(**X**). Now, the resulting row vector **v**^*(i)*^ adds up to 0. The reason [19] adopt this transformation is because the estimator of Cov[clr(**Y**)] can serve as a good approximation of Cov[clr(**X**)], which hence can be used for inferring the network structure.

While SEcov does not encode the graph using the covariance matrix or precision matrix, we still apply the clr transformation to reduce the potential experimental bias. As for the zero counts in the original count data **X**, we add a unit pseudo count before taking the log to avoid producing infinite values. Then we apply SEcov on the transformed data **v**^*(i)*^ to assess the association between any pairs of OTUs by (2).

### Microbial Network Selection by SEcov

Given a microbiome dataset with *d* species, there are *M* = *d*(*d* − 1)/2 pairs of OTU-OTU association in total to be evaluated. As described in Materials and Methods section, for each pair, one can use 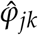 to assess the existence and strength of the association between OTUs. This implies that we need to simultaneously test *M* = *d*(*d* − 1)/2 null hypotheses. An edge between the *j*th and *k*th OTU is included in the recovered microbial network if and only if *H*_0*jk*_ is rejected, corresponding to *p*_*jk*_ ≤ *t* for *t >* 0. The threshold *t* is a pre-specified significance level, which should be appropriately selected especially for such a large-scale test. To this end, here we consider a multiple testing procedure by adjusting the significance level based on the sparsity of the graph, such that the false discovery rate (FDR) is well controlled.

So, in total there are *M p*-values, *{p*_*jk*_*}*_1≤*j<k*≤*d*_, smaller *p*-values provide stronger statistical evidence that *φ*_*jk*_ differs from zero. According to this rationale, we order all *p*-values,

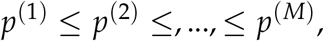

and reject those hypotheses having

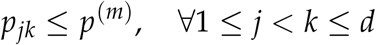

where *m* ∈ *{*1, …, *M}* is an integer. In this way, the total number of rejections is

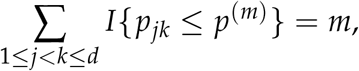

and the final network would include exactly *m* most significant edges/associations. The false discovery rate [30] can be defined by:

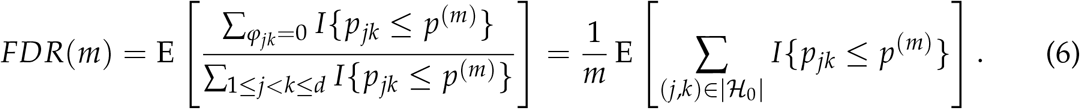

An important fact here is that *p*_*jk*_ should be uniformly distributed under the null hypothesis since the test statistic converges to a standard normal variable under *φ*_*jk*_ = 0. As a result, the FDR can be written as:

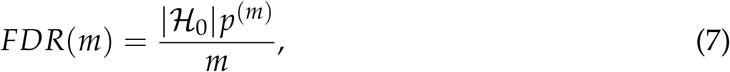

where |ℋ_0_| is the number of true null (i.e., no association). The estimate in (7) is a Benjamini–Hochberg-type FDR estimate that relies on the null *p*-values being approximately uniform, which follows from the asymptotic normality in (3).

In the context of microbial network analysis, the underlying graph is often sparse, say the number of true edges is roughly of *c* × *d* where *c* is a positive constant. Then, |ℋ_0_| can be approximated by (*d*^2^− *d*)/2 [23]. If we let *p*^*(m)*^ = *p*^*(d)*^ such that exactly *d* edges are included, then *FDR*(*d*) *≈* 0.5(*d* − 1) × *p*^*(d)*^, which is small whenever *p*^*(d)*^is small. Note that selecting exactly *d* edges is a sparsity-matching device; in practice one would instead fix a target level *α* and select the largest *m* with *FDR*(*m*) ≤ *α*.

In Figure **1**, we illustrate the procedure of network selection by tuning *p*-values in SEcov and compare it with other popular graphical modeling methods for microbiome data. The example microbiome data called American Gut Project [24] in an R package SpiecEasi [18] was used to generate a synthetic data using “Normal to Anything” (NorTA) approach [19]. The synthetic data includes 1000 samples and 127 OTUs, and is based on the cluster graph type and zero-inflated negative binomial distribution. The true number of edges equals to *d* = 127. The red and green color respectively represent the negative and positive association estimated by the method. Here. we consider three other methods, SpiecEasi [19], SPRING [36], and SparCC [12]. The former two models infer the conditional dependency by partial correlation, following the neighborhood selection methodology [26], and select the sparse graph by StARS model selection approach [22]. The third one measures the direct OTU-OTU association and edges in the graph are selected by a prespecified cutoff point.

**FIG 1.**
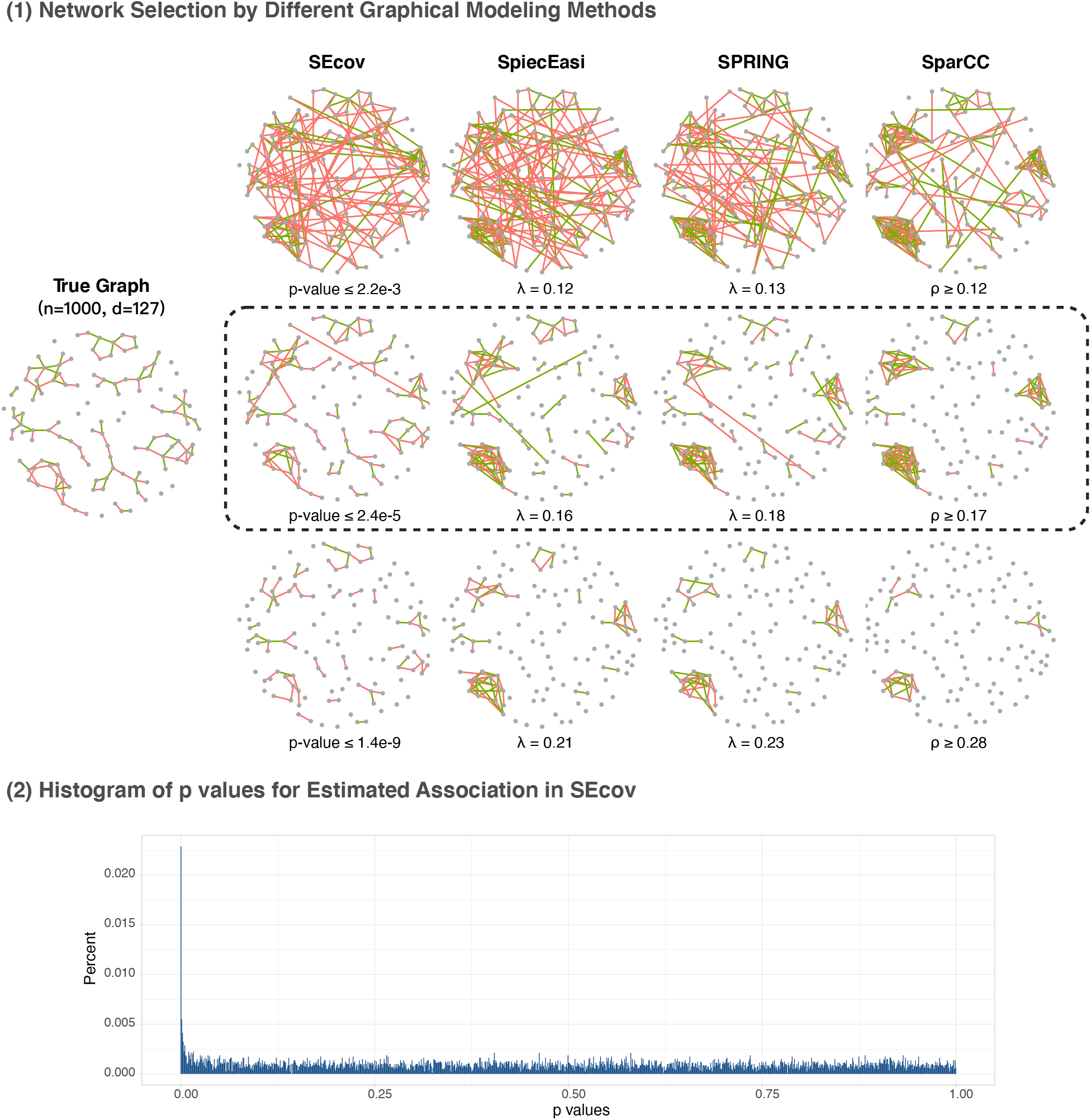
(1) Illustration of network selection for different graphical modeling methods and (2) the distribution of *p*-values for the estimated associations of all involved pairs of OTUs by SEcov.

For SEcov, edges are included if *p*-value is ≤ *t*. A smaller significance level *t* creates sparser networks. Correspondingly, SpiecEasi and SPRING derive a sparse graphical model by neighborhood selection principle where a larger regularization parameter *λ* results in a higher sparsity. As for SparCC, only those edges with correlation *φ* greater than a cutoff are included. The figures in the top line show that, when the estimated graph is too dense, more edges are incorrectly included and the networks are messy. On the other hand, when the graph is too sparse, see the bottom line of part (1), many true associations cannot be detected. However, those included edges reveal the strongest signal of association. For the second line within the dashed box, we choose the significance level in SEcov and correlation cutoff in SparCC such that exactly *d* = 127 edges are included for comparability across methods. For SpiecEasi and SPRING, we choose *λ* such that the number of detected edge approaches to 127 at most. In this case, the sparsity of the network is correctly characterized, with all four methods achieving their best performance among the cut-offs considered. Comparing the graphs, SEcov can best recover the topology of the underlying true graph with correct sign of the dependency.

Additionally, we also demonstrate the distribution of all *p*-values obtained by SEcov. If all associations were null, the *p*-values should follow a uniform distribution. However, the histogram (2) in Figure **1** indicates that a small proportion of 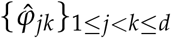 have extremely small *p*-values that violate uniformity, indicating that the corresponding null hypotheses should be rejected. If we select the significance level by *p*^*(m)*^= *p*^*(d)*^= 2.4 × 10^−5^ in this case, then *FDR*(*d*) *≈* 0.5(127 − 1) × 2.4 × 10^−5^ = 0.0015, which is controlled.

The demonstration in Figure **1** provides the first sign of the superiority and validity of SEcov for inferring microbial network. In next section, we shall conduct an overall synthetic study to compare the performance of these four methods.

## RESULTS

In this section, we conduct experiments to evaluate the performance of SEcov for inferring microbial networks from microbiome data and compare it with other popular graphical modeling approaches in the context of microbial association studies.

### Synthetic microbiome data generation

We follow the simulation setting used in [19] to generate the synthetic quantitative microbial abundance data, which are modeled on American Gut Project (AGP) data. This benchmark defines the underlying graph through a latent Gaussian precision-matrix construction: non-zero entries of the precision matrix encode the adjacency matrix of the underlying undirected graph, and the association strength is summarized by the corresponding partial correlation.

To simulate the data, we first construct a correlation matrix Σ. This can be done by specifying the properties of the precision matrix. Assume that there are *d* OTUs. We generate an undirected graph in the form of an *d* × *d* adjacency matrix Ω, where (*j, k*)-entry *ω*_*jk*_ = 1 if and only if there is an edge connecting the *j*th and *k*th node, otherwise *ω*_*jk*_ = 0. Here, we consider three graph types for the topology of the underlying networks: (i) band graphs; (ii) cluster graphs; and (iii) scale-free graphs, as shown in Figure **2**. These three types are representative and have been widely used for synthetic studies of microbial network analysis [16, 19, 36]. To control the sparsity of the graph, we let the total number of edges *e*, be equal to the number of nodes, i.e., *e* = *d*, for all networks. This corresponds to a sparse-network regime with average degree 2*e*/*d* = 2. The choice *e* = *d* fixes the sparsity of the true graph for benchmarking and is separate from the ROC/AUC analyses, which evaluate the full edge-ranking behavior of each method. We then convert Ω to a positive definite precision matrix Θ using an R package SpiecEasi [18]. The non-zero entries *θ*_*jk*_ *≠* 0 in Θ correspond to *ω*_*jk*_ = 1 in Ω for any 1 ≤ *j < k* ≤ *d*, which are uniformly sampled from [−3, −2] *∪* [2, 3]. Then we let *θ*_*kj*_ = *θ*_*jk*_ by the symmetry of the precision matrix. The diagonal entries of Θ are set to some constant to ensure that Θ is positive definite with a default condition number *κ* = 100. Finally, we obtain the correlation matrix Σ by taking the inverse of Θ, followed by scaling. Finally, “Normal to Anything” (NorTA) approach [5, 13] is applied to generate synthetic abundance data from this latent correlation structure, with marginal distributions specified as zero-inflated negative binomial distributions. Thus, while the benchmark graph is defined through a latent Gaussian precision matrix, the observed synthetic abundance data have non-Gaussian count marginals.

**FIG 2.**
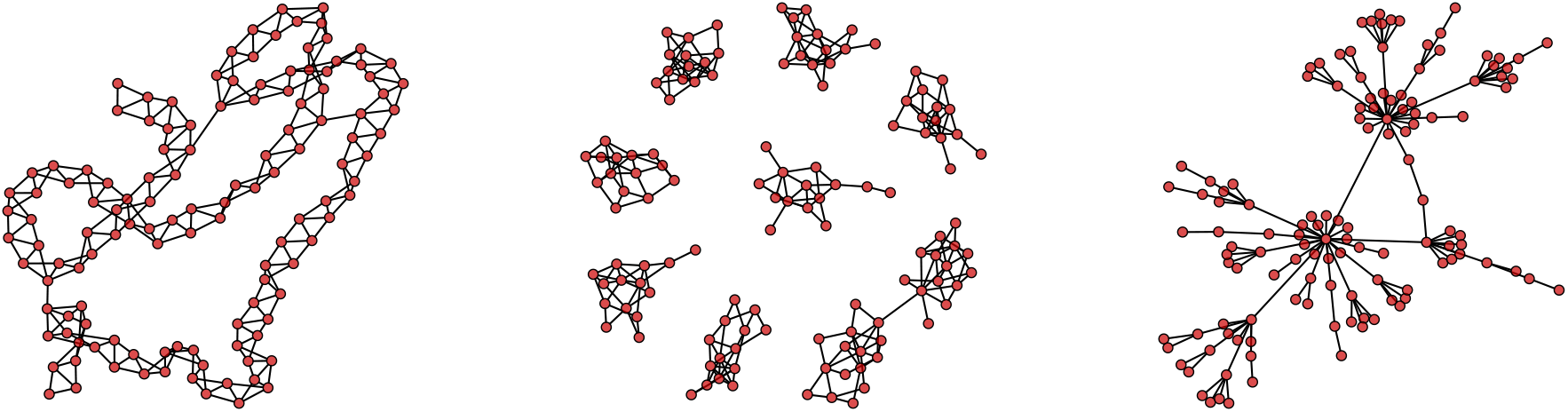
Three graph types considered for the synthetic microbiome data, band (left panel), cluster (middle panel), and scale-free graphs (right panel).

The AGP count data we use can be downloaded from https://github.com/zdk123/SpiecEasi/tree/master/inst/extdata, which contains 3738 OTUs across 407 samples. We first apply the same filtering steps as used in [19]: (1) remove the samples whose total sequencing depth falls below the first quartile of all sequencing depths (16,134 sequence reads) such that the data finally contains *n* = 305 samples; and (2) remove OTUs if the prevalence (present) is fewer than *γ*_1_ = 0.37 and *γ*_2_ = 0.60. Respectively, this leads to two filtered data sets: (1) a high-dimensional data with *d*_1_ = 192 OTUs; and (2) a low-dimensional data with only *d*_2_ = 68 OTUs. Obviously, more zero counts are included in the high-dimensional data. To assess the effects of sample size on network recovery, for both *d*_1_ and *d*_2_, we compare the performance of four graphical modeling methods on different sample sizes, *n* = 102, 300, 600, 1500.

Given synthetic abundance data with a known underlying structure, we can obtain the true partial correlation matrix *P* by standardizing the precision matrix and changing the sign of the off-diagonal entries. This enables benchmarking of different sparse inverse and partial correlation estimation techniques, where *ρ*_*jk*_ in *P* will serve as the ground truth of the strength of conditional association. We use “wi2net” function in an R package qgraph [8] to turn a precision matrix into the partial correlation matrix. We then apply SEcov to recover the microbial network as described in Materials and Methods section: (i) apply the clr-transformation to the abundance data; (ii) calculate 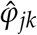 by (2) where the conditional means are estimated by random forest algorithm (implemented by “ranger” function in an R package ranger [34]), and obtain p-values *p*_*jk*_ for all 1 ≤ *j < k* ≤ *d*; (iii) rank all taxon pairs by their *p*-values. For selected-network summaries of SEcov, we select the *m* pairs with the smallest p-values, equivalently thresholding at *p*^*(m)*^. In the sparsity-matched selected-network summaries, we set *m* = *e* = *d*, matching the known true network size in the synthetic generator. This selected-network summary is distinct from the ROC/AUC analyses, which use the complete ranking of all taxon pairs. The selected edge weights are given by the corresponding SEcov estimates 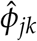, which are compared with the true partial correlations *ρ*_*jk*_.

For comparison, we consider another three popular graphical modeling approaches for microbiome data: (1) SpiecEasi’s method [19]; (2) SPRING’s semi-parametric rank-based correlation estimation [36]; and (3) SparCC [12], also designed to be robust to compositional artifacts. SpiecEasi and SPRING (respectively implemented in R packages SpiecEasi and SPRING) infer the conditional dependency, following the neighborhood selection methodology [26] to derive a sparse graph where the estimated coefficient matrix 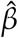 is symmetrized by taking the maximum absolute value, i.e., 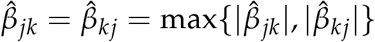 . For both of them, the regularization parameter *λ* is tuned by StARS model selection approach [22]. The edge weights *ρ*_*jk*_ of SpiecEasi and SPRING are respectively estimated by the output coefficient 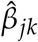 based on the connection between partial correlation and regression coefficients [11]. Since [36] recommend directly applying SPRING on the absolute abundance data for a better performance, here we do not transform the microbiome count data when using SPRING method. SparCC (also implemented in an R package SpiecEasi) infers the marginal association with the sample correlation 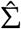, where edges having 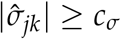 are selected (*c*_*σ*_ *>* 0 is a prespecified cutoff) [12]. The edge weights of SparCC inferred network are the sample correlation 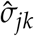 . Table **1** summarizes the four methods for making comparison in all of our experiments.

**TABLE 1.**
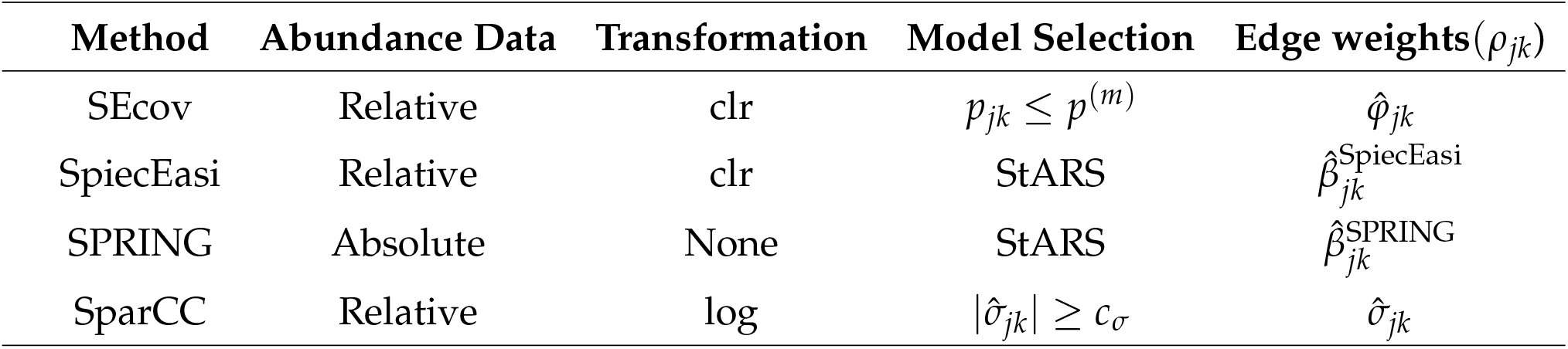
Summary of four methods for comparison.

### Simulation results

To evaluate the overall performance of each method to infer the underlying networks, we show the receiver operating characteristic (ROC) curve for all scenarios. In ROC curves, the y axis is the true positive rate (TPR) and the x axis is the false positive rate (FPR):

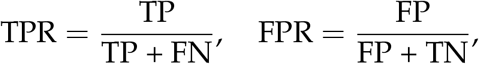

where TP, TN, FP, FN represent the number of true positive, true negative, false positive, and false negative results. For SEcov, edge predictions are ranked by *p*-values. For SpiecEasi and SPRING, ROC curves are traced along their regularization paths. SparCC edge predictions are ranked by the magnitude of estimated correlations. The results are provided in Figure **3**. The figure shows that the performance of all four methods improves in terms of ROC curve as sample size increases. Given the number of OTUs (*d*) and the sample size (*n*), networks with band-type structure are the easiest to recover while scale-free networks are most challenging for all methods. When the sample size is very small (*n* = 102), SEcov performs the best when the dimension is moderate *d* = 68. When *d* = 192, all four methods have similar performance with almost overlapping curves. As sample size increases, SEcov always exhibits better performance regardless of the dimension and underlying graph type.

**FIG 3.**
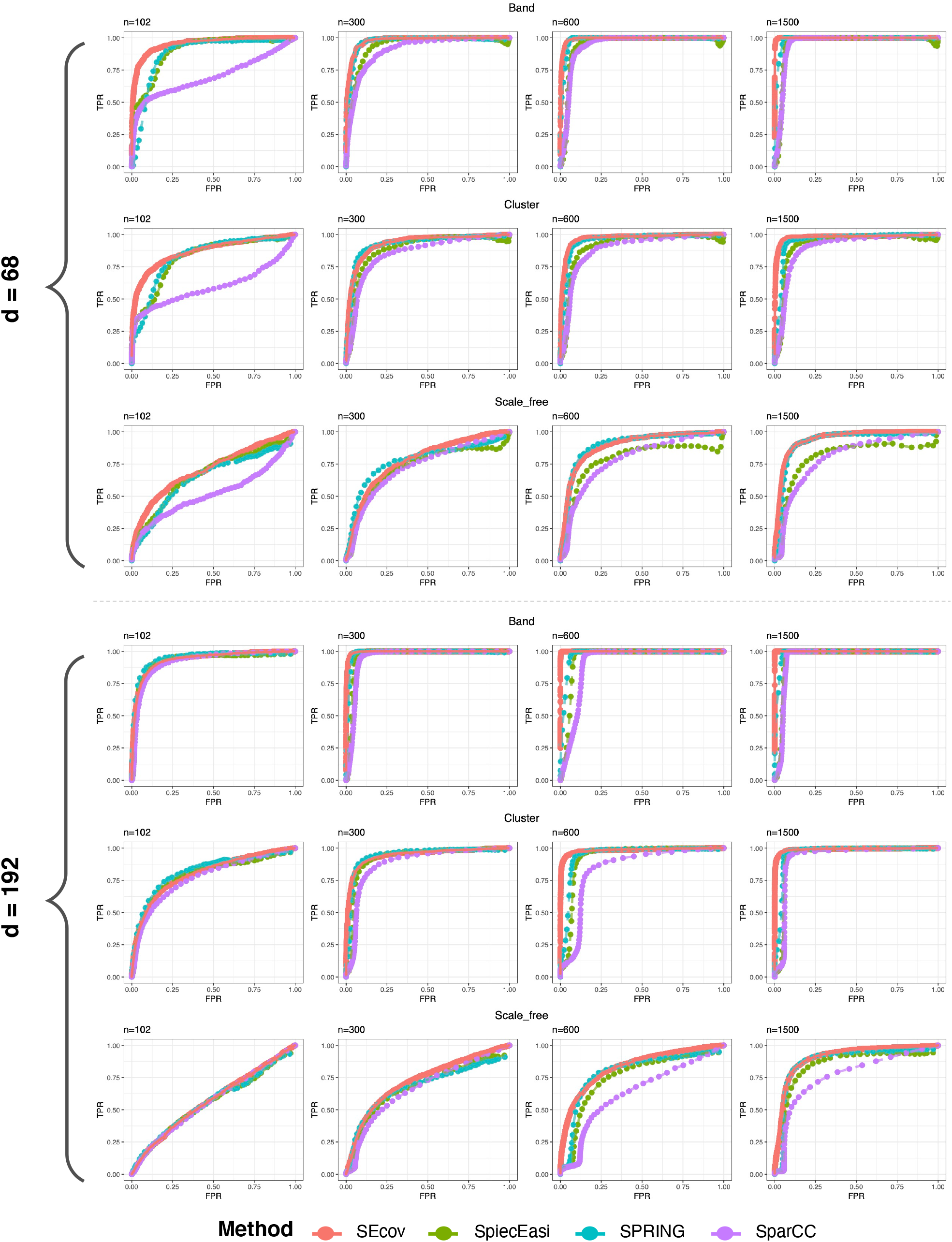
ROC curves of SEcov, SpiecEasi, SPRING, SparCC for 24 scenarios: 3 graph types (band, cluster, scale-free) x 2 dimensions (*d* = 68, 192) x 4 sample size *n* = 102, 300, 600, 1500. The curve is averaged over 10 synthetic datasets.

To accurately see the trend, we calculate the area under the ROC curve (AUC values), for two representative scenarios: (i) *n* = 1500, *d* = 68 (large sample with low dimension); and (ii) *n* = 600, *d* = 192 (moderate sample with high dimension). The results are shown in Table **2**. In both settings, SEcov always achieves the best performance among these methods. It can achieve near-perfect recovery with AUC value close to 1 when the sample size is 1500 and graph is band-type or cluster-type. Because these ROC/AUC comparisons are based on full rankings or regularization paths, they do not depend on the final choice of selecting exactly *d* edges for SEcov.

**TABLE 2.** AUC values (standard deviation) of SEcov, SpiecEasi, SPRING, SparCC for 3 graph types under (i) *n* = 1500, *d* = 68 and (ii) *n* = 600, *d* = 192.

| Dimension | Graph | SEcov | SpiecEasi | SPRING | SparCC |
| --- | --- | --- | --- | --- | --- |
| n=1500, d=68 | Band | 0.999 (0.000) | 0.954 (0.002) | 0.957 (0.008) | 0.955 (0.003) |
|  | Cluster | 0.980 (0.011) | 0.923 (0.009) | 0.923 (0.014) | 0.905 (0.013) |
|  | Scale-free | 0.939 (0.011) | 0.800 (0.040) | 0.886 (0.023) | 0.804 (0.024) |
| n=600, d=192 | Band | 1.000 (0.000) | 0.915 (0.008) | 0.950 (0.006) | 0.901 (0.012) |
|  | Cluster | 0.984 (0.006) | 0.893 (0.007) | 0.916 (0.010) | 0.841 (0.013) |
|  | Scale-free | 0.832 (0.036) | 0.722 (0.033) | 0.754 (0.046) | 0.636 (0.029) |

To see the best performance each method can potentially achieve, we also calculate the Hamming distance, which is defined by the total number of edge disagreements between the recovered network and the ground truth, i.e., Σ_1≤*j<k*≤*d*_|*ê*_*jk*_ − *e*_*jk*_|. Given the number of OTUs *d*, the Hamming distance can range from 0 to *d*(*d* − 1)/2. Since the four graphical modeling methods use different threshold methods, here we only plot the range of thresholds of each method where the minimum Hamming distance can be reached. The mean Hamming distance values over 10 replications as function of different thresholds are plotted in Figure **4**. For both band-type and cluster-type graphs, SEcov outperforms all the other methods. The minimum Hamming distance reveals that SEcov can have a nearly perfect edge prediction if *p*^*(m)*^is appropriately selected when the sample size is large enough. In fact, under these scenarios, *p*^*(m)*^= *p*^*(d)*^mostly approaches the lower bound. SPRING has the secondary best performance for these two graph types, which however is not comparable to SEcov. Networks with scale-free topology are the hardest to infer: all four methods have comparable performance in terms of the minimum edge disagreement.

**FIG 4.**
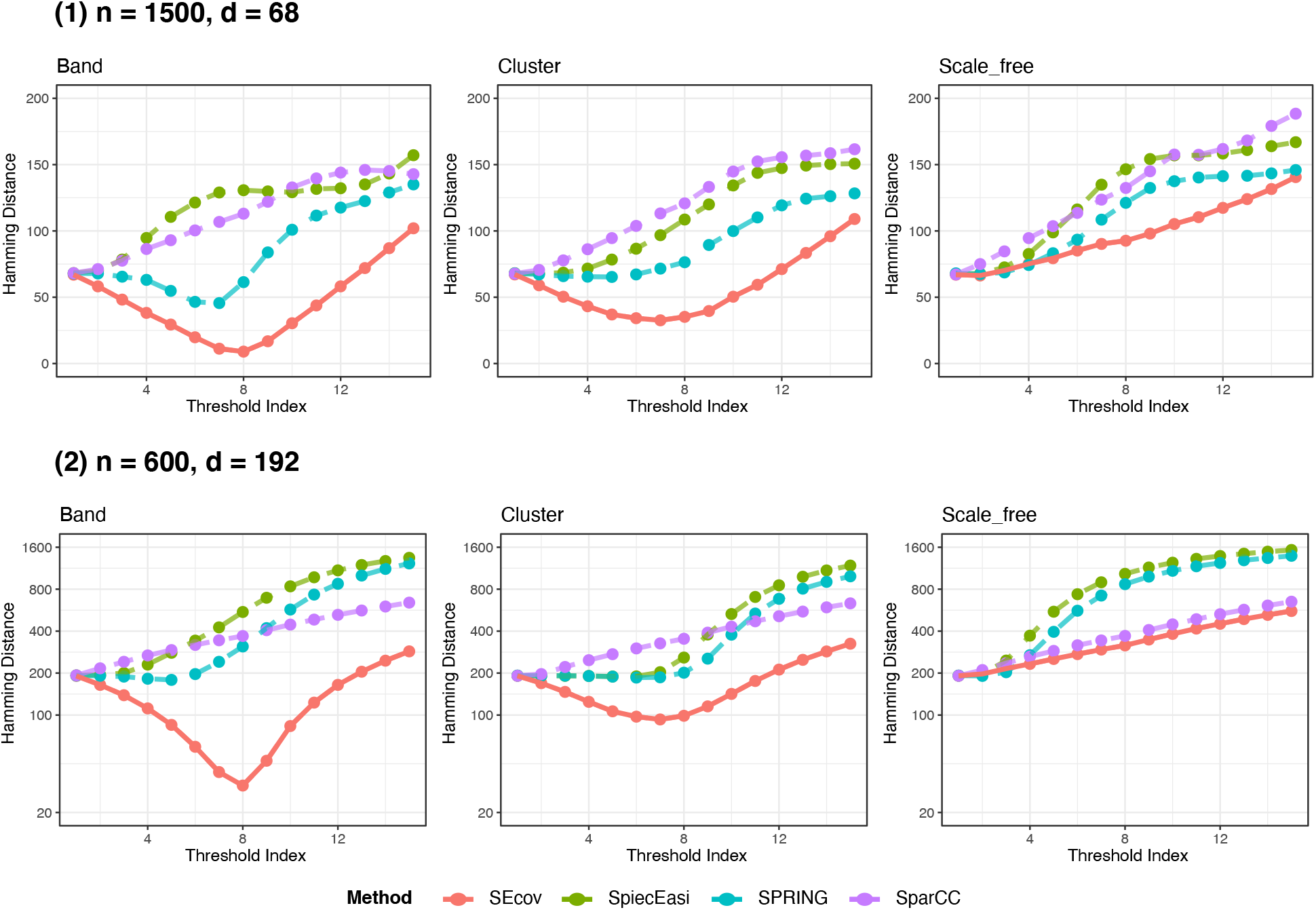
Hamming distance for different types of graph estimated by four graphical modeling methods. For SEcov, SpiecEasi, SPRING, and SparCC, the mean Hamming distance values are respectively plotted as the function of (i) different significance level *p*^*(m)*^∈ (*p*^(1)^, *p*^(170)^); (ii) different values of the regularization parameter *λ* ∈ (0.4, 0.1); (iii) different values of the regularization parameter *λ* ∈ (0.6, 0.16); and (iv) different cutoff values in (0.35, 0.1). The mean Hamming distance values are calculated over 10 replications.

We next consider the discrepancy between the final recovered networks and the ground truth. To this end, the network from SEcov is selected by 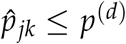 such that the most significant *d* edges are included. For SpiecEasi and SPRING, the optimal graphs are both selected by StARS approach with stability threshold *t*_*s*_ = 0.05. SparCC includes edges with high correlation: 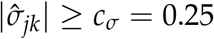 . These final-network summaries compare the methods at their method-specific operating points: top-*d p*-value selection for SEcov, StARS-selected graphs for SpiecEasi and SPRING, and a fixed correlation cutoff for SparCC.

We first plot the false discovery proportion (FDP) and the mean absolute error (MAE) which is

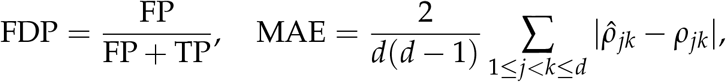

in Figure **5**. The FDP is the realized proportion of false edges in a single dataset. For SEcov, its average over replicates provides an empirical counterpart to the FDR targeted by (6)–(7); for the other methods, FDP is used as a descriptive measure of false-edge burden at their selected operating points. At these method-specific operating points, SEcov shows the lowest FDP and MAE in most scenarios. Particularly, one can observe the trend that FDP is decreasing with increasing sample size. This is consistent with the asymptotic control in (7): as *n* grows relative to *d*, the null p-values approach uniformity and the realized FDP stabilizes. When the sample size is large (*n* = 1500), the FDP for SEcov is low (≤ 0.2) for both band- and cluster-type graphs regardless of dimension, whereas the other methods do not exhibit this behavior. Unfortunately, other methods are unable to display these properties. In fact, the relationship between the regularized parameter and the number of false edges in SpiecEasi/SPRING estimation is unclear. As for SparCC, it fixes the threshold by *c*_*σ*_ = 0.25, which is not directly tied to FDP. Thus, because these methods are not tuned through a *p*-value, they offer no mechanism to target the false discovery rate and cannot guarantee a low risk of inferring spurious associations. For MAE, we observe similar patterns: the distance between the SEcov-inferred networks and the ground truth goes down as sample size increases for all graph types, which cannot be observed in other estimation methods. In fact, the edge weight estimation 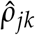 in SpiecEasi/SPRING are assigned by the regression coefficients 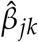 from neighborhood-selection methods [26], which however do not explicitly estimate the partial correlation. Sample correlation used by SparCC for the edge weights is also different from partial correlation.

**FIG 5.**
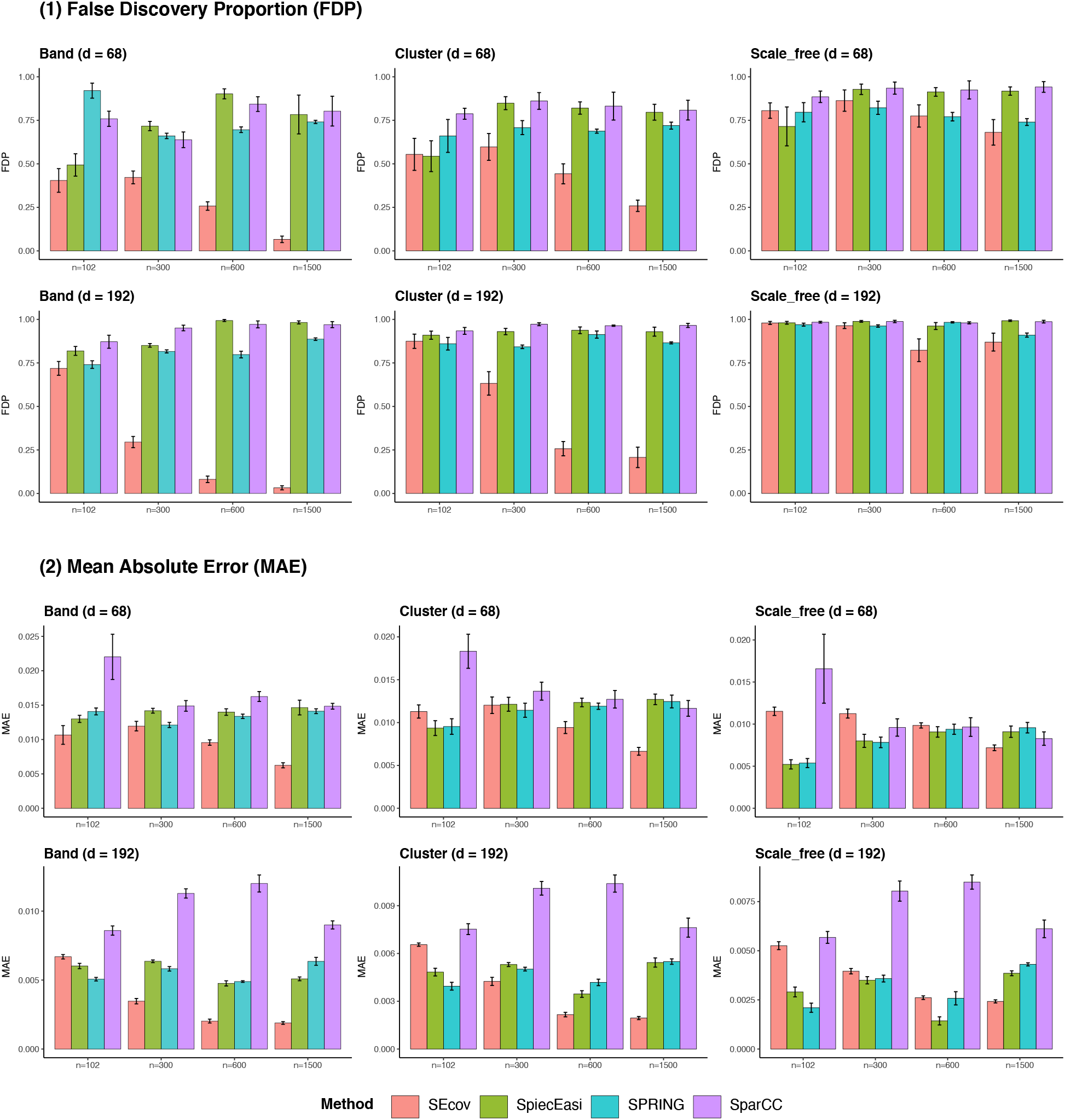
Barplots of false discovery proportion and mean absolute error for four graph estimation methods under all synthetic scenarios. The height of the bar represents the average values across 10 replicates.

Finally, we focus on the cluster-type graph and analyze the overlapping sets of selected edges in the two settings again. The experiments are repeated 10 times and we calculate the mean values as well as the standard deviation in the parentheses across 10 replicates. The Venn diagram in Figure **6** shows: (i) the average number of edges that overlap across the methods indicated by the overlapping area; and (ii) the proportion of correctly inferred edges of all overlapping edges in each area. When *n* = 1500, *d* = 68, almost all edges detected by SEcov can be identified by other methods. However, for other sets uniquely/pairwisely identified by methods without SEcov, the true positive proportions are extremely low. Only those sets with SEcov included are likely to have a high proportion of true positive. The highest true positive rate (TPR), 0.98, happens in the set, where edges are only jointly identified by SEcov and SPRING. This is followed by the set where edges are jointly selected by SEcov, SPRING and SpiecEasi, TPR is 0.73. When *n* = 600, *d* = 192, the set with the highest TPR, 0.86, is also the one with edges that are only selected by SEcov and SPRING with around 45 edges included in this set. The second highest TPR happens in the set uniquely identified by SEcov, which is 0.84 of 100 edges. In both low- and high-dimensional cases, the Venn diagram suggests not considering the statistical associations that are uniquely identified by SPRING, SpiecEasi, SparCC due to the low TPR. Instead, it is more promising to select the edge sets of SEcov that also overlap with other approaches, such as SPRING and SpiecEasi.

**FIG 6.**
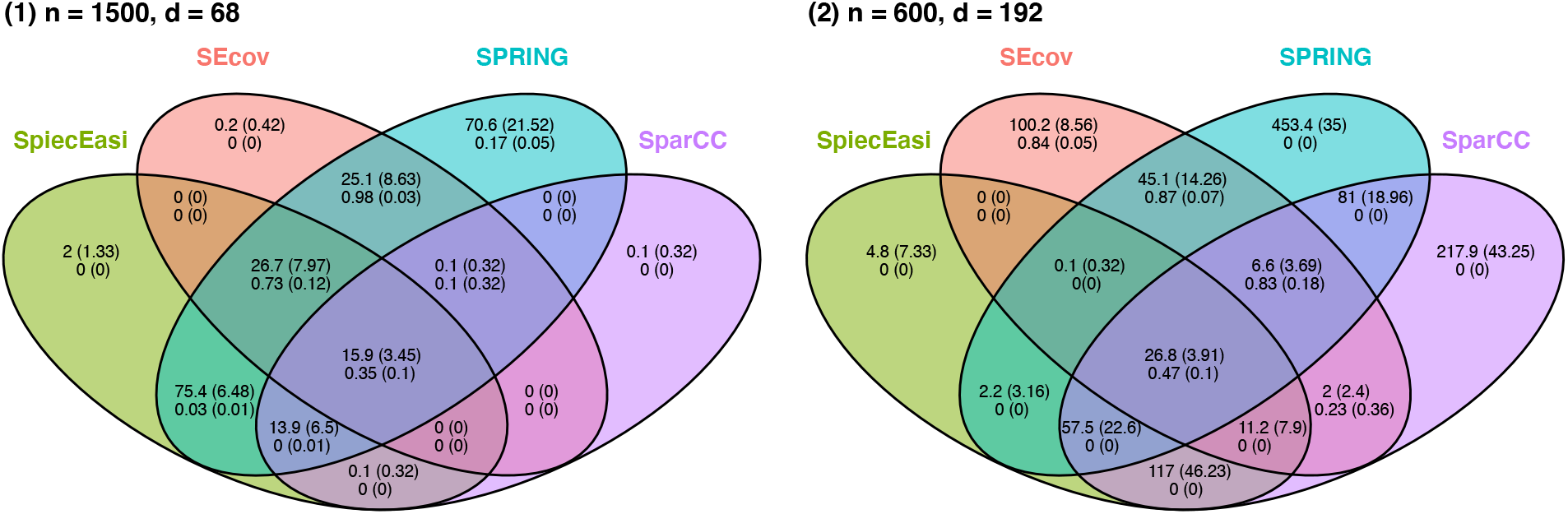
Average number of edges (top row) and average proportion of correctly identified edges in each overlapping set (standard deviation) indicated by the Venn diagram. The mean values are calculated across 10 replicates.

### Analysis of the PIN Cohort

We apply SEcov to infer microbial association networks using vaginal microbiome data collected during the third trimester of pregnancy in the PIN cohort. The data analyzed here are species-level Stirrups count profiles. Stirrups is a vaginal microbiome-specific species-level classification approach developed for 16S rDNA reads using a curated vaginal 16S rDNA reference database [10, 31].

The original Stirrups count table contained 910 samples and 460 taxa. Before network construction, we apply filtering to reduce the influence of rare taxa and low-depth samples. Specifically, taxa present in fewer than 10% of samples are removed, and samples with total sequencing depth below the first quartile of the sequencing-depth distribution are excluded. This filtering results in a final analytic table with *n* = 682 samples and *d* = 73 taxa. To account for the compositional nature of microbiome count data and avoid undefined log-ratio values from zero counts, we add a unit pseudocount and applied the centered log-ratio transformation before estimating microbial associations.

SEcov is then applied to all *d*(*d* − 1)/2 = 2, 628 taxon pairs. For each pair of taxa, the conditional means are estimated using random forests with 1,000 trees, and the SEcov statistic and its corresponding two-sided *p*-value are computed using the asymptotic normal approximation. The final SEcov network is obtained by selecting the *d* = 73 most significant taxon pairs, following the same sparsity-matched selection strategy used in the simulation study.

For comparison, we also construct microbial networks using SpiecEasi, SPRING, and SparCC. SpiecEasi and SPRING are used as conditional-dependence network methods, with regularization selected by the StARS procedure, whereas SparCC is used as a compositionally robust marginal correlation method. For SparCC, the *d* = 73 taxon pairs with the largest absolute correlations are selected so that its network size was comparable to that of SEcov. The resulting networks are compared in terms of edge overlap and sign agreement.

Figure **7** shows the inferred networks by SEcov as well as the number of overlapping edges by four graphical modeling methods. Table **3** summarizes the agreement of edge signs inferred by different graphical models. SEcov selects 73 associations, of which 49 are positive and 24 are negative. In contrast, the networks inferred by SpiecEasi, SPRING, and SparCC are dominated by positive associations: 92%, 100%, and 97% of their selected edges, respectively, are positive. Notably, the 24 negative associations identified by SEcov are almost entirely missed by the competing methods. None of these negative SEcov edges are selected by SpiecEasi or SPRING, and only one is also selected as negative by SparCC. By contrast, positive SEcov edges show greater overlap with the other methods: 28, 27, and 15 positive SEcov edges are also selected by SpiecEasi, SPRING, and SparCC, respectively.

**TABLE 3.** Two-way tables, summarizing the agreement of the sign of edges inferred by different graphical model methods.

|  | SpiecEasi |  |  | SPRING |  |  | SparCC |  |  |
| --- | --- | --- | --- | --- | --- | --- | --- | --- | --- |
|  | – | 0 | + | – | 0 | + | – | 0 | + |
| SEcov |  |  |  |  |  |  |  |  |  |
| – | 0 | 24 | 0 | 0 | 24 | 0 | 1 | 23 | 0 |
| 0 | 5 | 2518 | 32 | 0 | 2517 | 38 | 1 | 2498 | 56 |
| + | 0 | 21 | 28 | 0 | 22 | 27 | 0 | 34 | 15 |

**FIG 7.**
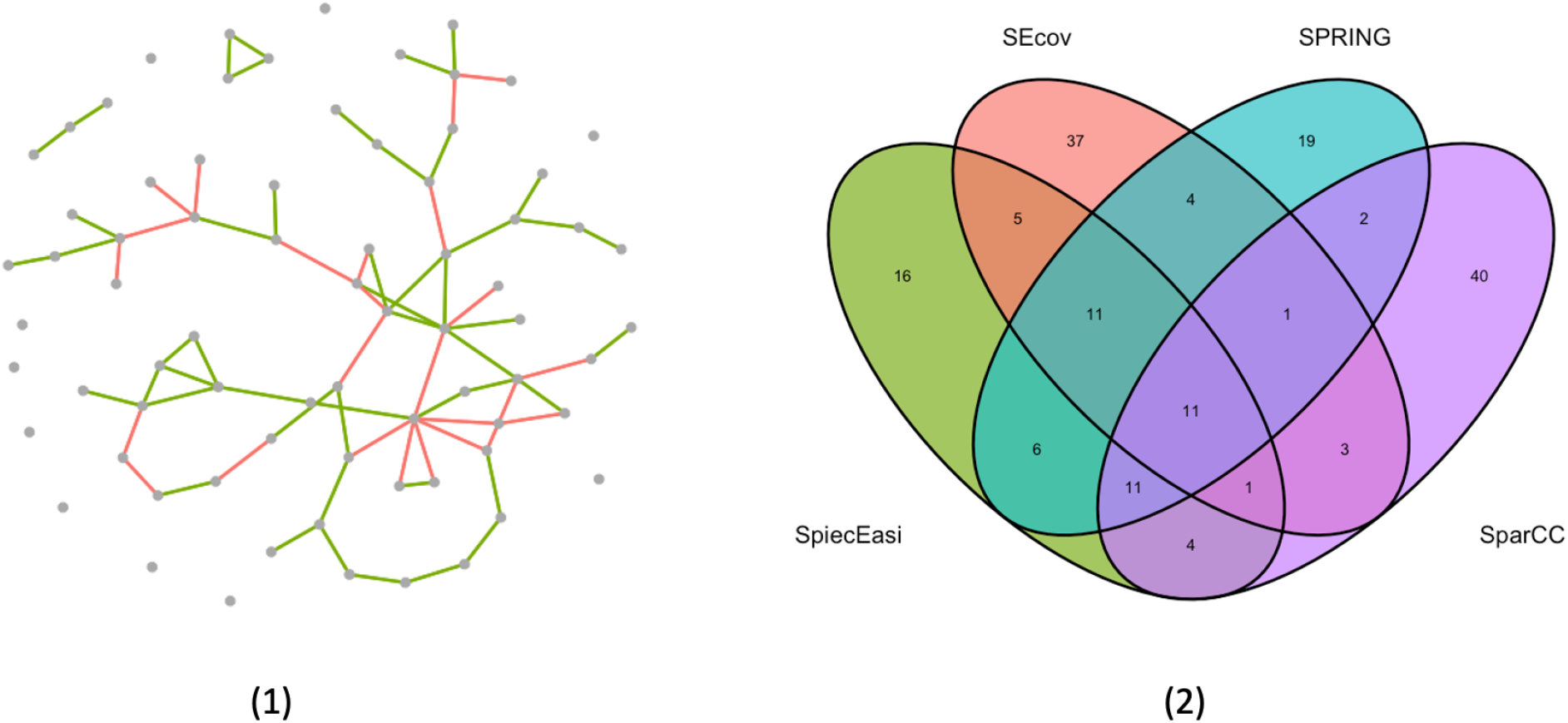
Results of Stirrups data analyses. (1) The recovered network by SEcov. The corresponding network with taxon labels is provided in Supplementary Fig. S1; (2) The number of overlapping edges by Venn Diagram of four different association networks.

These results suggest that positive microbial associations are more consistently recovered across methods, whereas negative conditional associations are more sensitive to the modeling approach. Because SEcov adjusts flexibly for the remaining taxa through nonparametric conditional-mean estimation, it may reveal conditional mutual-exclusion patterns that are not captured by methods relying primarily on sparse linear graphical modeling or marginal correlation. Nevertheless, these edges should be interpreted as conditional statistical associations rather than direct ecological or causal interactions.

## DISCUSSION

We proposed a practical framework for constructing microbiome co-occurrence networks based on SEcov, a nonparametric measure of conditional association. By adjusting for the remaining taxa through conditional-mean functions, SEcov can accommodate nonlinear relationships and avoids the strong Gaussian and linearity assumptions required by many graphical-modeling approaches. In addition, its asymptotic normality provides a basis for edge-level inference, thereby improving the interpretability of the resulting network compared with methods that rely primarily on correlation cutoffs or regularization tuning parameters. Through simulation studies, we showed that SEcov can improve network recovery across multiple graph topologies, dimensions, and sample sizes. The synthetic benchmarks used here follow a standard NorTA/precision-matrix construction, which provides a known graph structure and non-Gaussian count marginals for comparison with existing microbial network methods. More general data-generating mechanisms with explicitly nonlinear conditional dependence remain an important direction for future simulation studies.

To utilize SEcov for evaluating the conditional dependence in practice, we need to calculate the conditional means and we adopt the random forest algorithm to estimate the conditional means. However, this technique is flexible and other machine learning algorithms can be used to estimate the conditional means, such as generalized additive models, locally weighted linear regression, and LASSO [33]. The choice of a suitable machine learning algorithm would enhance the performance of the new approach and domain knowledge should be used to design the algorithm that is sensitive to the signal of interest.

When it comes to the context of microbial interaction, normalization is a critical step as the abundances or counts of microbial species typically vary in magnitude and encounter issues such as zero-inflation or over-dispersion. In this paper, we propose to use the centered log-ratio transformation for the compositionality before conducting network construction since it can serve as a nice approximation to infer the network structure. Other normalization methods can be used, such as rarefaction [25], total sum scaling [7], and cumulative sum scaling [27]. Each methods has its own advantages, but some concerns are discussed in the literature (see [32]). To bypass the challenge of picking an optimal normalization strategy, one can consider an omnibus approach to obtain the value of SEcov [29].

## ACKNOWLEDGMENTS

This work is supported, in part, by the National Institutes of Health grant R01GM129512 and the National Research Foundation of Korea(NRF) grant funded by the Korea government(MSIT) (RS-2022-NR068758, RS-2026-25471551).

## DATA AVAILABILITY STATEMENT

The codes for implementing the proposed method are available on GitHub at https://github.com/hoseungs/mNetwork. The PIN microbiome count data and analysis scripts used for the published PIN microbiome study are publicly available through the PINmicrobiome repository, and sequencing data are available through NCBI BioProject PRJNA694098.

## Supplemental Material

Supplemental Material includes the expression of estimator of 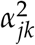 and the recovered network of Stirrups data with node names.

## Notes

### Competing Interest Statement

The authors have declared no competing interest.

